# PID1 alters antilipolytic action of insulin and increases lipolysis via Inhibited the activation of AKT/PKA Pathway

**DOI:** 10.1101/317024

**Authors:** Chunyan Yin, Yan Jin, Yuesheng Liu, Li Wang, Yanfeng Xiao

**Affiliations:** The Second Affiliated Hospital of Xi’an Jiaotong University, Xi’an, Shan Xi 710049, People’s Republic of China

**Keywords:** PID1, insulin, antilipolytic effect, lipolysis, AKT Pathway

## Abstract

**Purpose:** The aim was to investigate the mechanism for impaired control of lipolysis in obesity by investigating the effect of PID1 on insulin-induced activation of AKT/PKA/HSL pathway and lipolysis.

**Methods:** First, PID1 expression was detected in adipose tissue and blood insulin and glycerol levels were measured in high-fat diet-induced obese rats. Next, we examined the effect of different concentrations of insulin on lipolysis and AKT/PKA/HSL pathway in 3T3-L1cells. We also investigated the role of PID1 in regulating AKT/PKA/HSL cascade and lipolysis after insulin treatment and lipofectamine over-expression.

**Results:** PID1 expression is increased in adipose tissue from HFD rat and positive correlation with insulin levels and lipolysis. In 3T3-L1 adipocytes, we found that antilipolytic effect of insulin is mediated by AKT and AKT activated by insulin can results in phosphorylation of PKA and HSL and suppresses glycerol release. However, over-expression of PID1 counteracts insulin action as indicated by glycerol releaseand reduced level of Akt phosphorylation in accordance with a decrease in the activity of insulin-dependent PKA/HSLsignaling cascade.

**Conclusions:** All together, these data showed that activation of PID1 in adipose tissue increases lipolysis by altering the antilipolytic action of insulin. This suggests that PID1 may constitute a new strategy to ameliorate adipocyte lipolysis and hence to improve insulin sensitivity.

## Introduction

Obesity is an increasing global health problem that is usuallyaccompanied byinsulin resistance and type 2 diabetes. Elevated serum FFA levels are frequently present in obesityand there is substantial evidence implicating elevated freefatty acid levels was a consequence of inappropriate lipolysis asa major etiological factor for insulin resistance and type 2diabetes mellitus (T2DM) [1–2]. Thus, understandingin detail the mechanism by which the impaired insulin suppresses fat cell lipolysis is critical to identifying the underlying defect in resistantadipose tissue and ultimately developing effective therapeutics.

As it is well known, insulin, an important hormone for regulating glucose metabolism, is also a key hormone promoting lipogenesis and inhibiting lipolysis[3]. The antilipolytic effect of insulin has been proposed toinvolve the reduction of cAMP levels and thus PKA activity. In this model, insulin signaling activates phosphodiesterase3b (PDE3b) via the Akt-mediated phosphorylation of Ser273[4, 5]. The activationof PDE3B catalyzes the hydrolysis of cAMP and leads to lowering of the cellular levelof cAMP. Thelowering of cAMP further inhibits protein kinase A (PKA) activity and thereby a decrease hormone sensitive lipase and lipolysis [6], but recent results show that, PDE3B activities in obese patients were significantly reduced in adipose tissue [7]. Thus, decreased AKT/PDE3B pathway activity may be a contributing factor to the diminished antilipolytic effect of insulin in obese patients. PID1 were subtracted from normal-weight subjects using suppression subtractive hybridization (SSH)[8]. Guo et al found that PID1 which contain a phosphotyrosine binding (PTB) domain canbind to phosphorylated tyrosine residuesand impair insulin signal transduction. And increased expression of PID1 leads to a reduction in insulinstimulatedglucose uptake and impaired insulin-stimulated GLUT4translocation in mature adipocytes [9], but whether PID1 alsoinfluencessignaling pathway of insulin regulates lipid metabolism still needs to be confirmed by further investigations.

In this study, we examined the effects of PID1 on lipolysis in high-fat diet (HFD)–induced obese rats, and further investigated the potential molecular mechanism underlying these effects using 3T3-L1 cells. We present evidence that PID1 alters antilipolytic action of insulin by inhibiting the AKT/PKA pathway which was activated by insulin and lead to lipolysis in obese.

## Methods

### Animal care and treatments

Ninety-six and 3-wk-old male SD rats were individually housed in a humidity controlled room with 12 h light/dark cycle. All the rats consumed a commercial diet for 1 week. After that, animals were randomized into two dietary groups according to ratio of 1:2: chow (36, CH, 12% kcal fat) or high fat (72, HF, 60% kcal fat). Eight normal diet rats and sixteen fat diet rats were randomly selected and body weights were measured at the following time points: 8, 16, 20, and 24w. The experimental protocols were approved by the Animal Care and Protection Committee of Xi’an Jiao tong University.

### Blood chemistries

All rats were killed and samples of blood were collected to measure insulin and glycerol by ELISA (Sigma) at 8, 16, 20 and 24w. Enzymatic assay kits (Applygen)were used for the determination of serum glucose. Samples of adipose tissue were collected to detect PID1 immunohistochemistry.

### Immunohistochemistry

Rat adipose tissue were fixed with 4% paraformaldehyde and embedded in paraffin, and 5μm-sections were prepared. After paraffin removal, tissue sections were stained with insulin and PID1 antibody(Abcam).

### Cell culture

3T3-L1 were cultured in flasks (25 cm^2^) in phenol red-free Dulbecco’s modified Eagle’s medium. Differentiation was induced using described protocols [10]. When more than 90% cells were fully differentiated, cells were treated with varying insulin doses (1-100nmol/L). To block PKA pathway, the inhibitor of PKA (H-89) was treated 12 h after the exposure to 100nM insulin for 24h, then culture medium and cells were separated and stored.

### Lipolysis measurement

An aliquot of the media (400μl) was collected, and glycerol release in cell culture medium was determined by using a colorimetric method (Sigma). The amount of glycerol was normalized to protein concentration as an index of lipolysis.

### Immunoflourescence

3T3-L1 cells were cultured and differentiated on coverslips, then fixed with 4% paraformaldehyde for 20 min, permeabilized with 0.05% Triton X-100 in PBS (15min), and blocked with 5% BSA in PBST (1h at room temperature). Staining with PID1 antibody was followed by incubation with Alexafluor (488)-conjugated secondary antibodies(Jackson), then stained with 0.2μg/mL Nile Red (Sigma) for 5 min, and incubated with 0.1μg/mL DAPI for2 min.

### Immunoblot

Cells were lysed ice-cold RIPA buffer. After protein concentration had been measured, the samples were mixed with Laemmli sample buffer and subjected to polyacrylamide gel electrophoresis (PAGE)(10% acrylamide) and Western blot analysis. After the electro transfer of proteins onto a PVDF membrane (Millipore), membranes were incubated overnight at 4°C with continual motion, using specific primary antibodies (AKT, p-AKT, PKA, p-PKA, HSL, p-HSL). Detection of protein–antibody immune complexes was achieved using horseradish peroxidase-conjugated secondary antibodies diluted1:10000 in PBS with 0.05% Tween. The chemo luminescent signal was analyzed and quantified with use of the bio-rad system.

### PID1 over-expression construct and transfection

The coding sequence of PID1 was subcloned into thepcDNA3.1Myc/His Bvector to generate a plasmid expressing PID1 His fusion protein. Expression vectors carrying the PID1 coding sequence or empty vectors were transfected into differentiated 3T3-L1 using Lipofectamine 2000. Two days after transfection, 0.8 mg/ml G418 (Roche, Basel, Switzerland) was added to the medium to select for transfected cells. Drug-resistant cells began to form small colonies after two weeks of G418 addition. Individual colonies were isolated, propagated and PID1 was identified by RT-PCR.

### RNAi

Differentiated adipocytes were transfected with 30 nM siRNA targeting AKT with Turbofect (Thermo Scientific) according to the manufacturer’s instructions. The knockdown efficiency was evaluated by RT-PCR and Western blot analysis.

### Statistical analysis

Normal distributions were assessed by the Kolmogorov–Smirnov test. Results are expressed as means±SE. Comparisons between groups were assessed using t-test or one-way ANOVA with post hoc Bonferroni corrections as appropriate. Differences were considered significant at P <0.05.

## Results

### Changes of body weights and adipose tissue weights

The body weights (BWs) for high fat diet (HFD) rats and normal diet (ND) rats were measured weekly. Initial mean BWs of the 2groups did not differ significantly. At week 8, HFD groups had significantly higher BWs than ND groups, and the BWs remained higher throughout the 24-week dietary period(Figure. 1a). Similarly, after 8 weeks, epididymal and perirenal fat depot weights in HFD groups were heavier than that of ND groups (Figure. 1b), reflecting the high fat diet increased body fat content.

**Figure 1.**
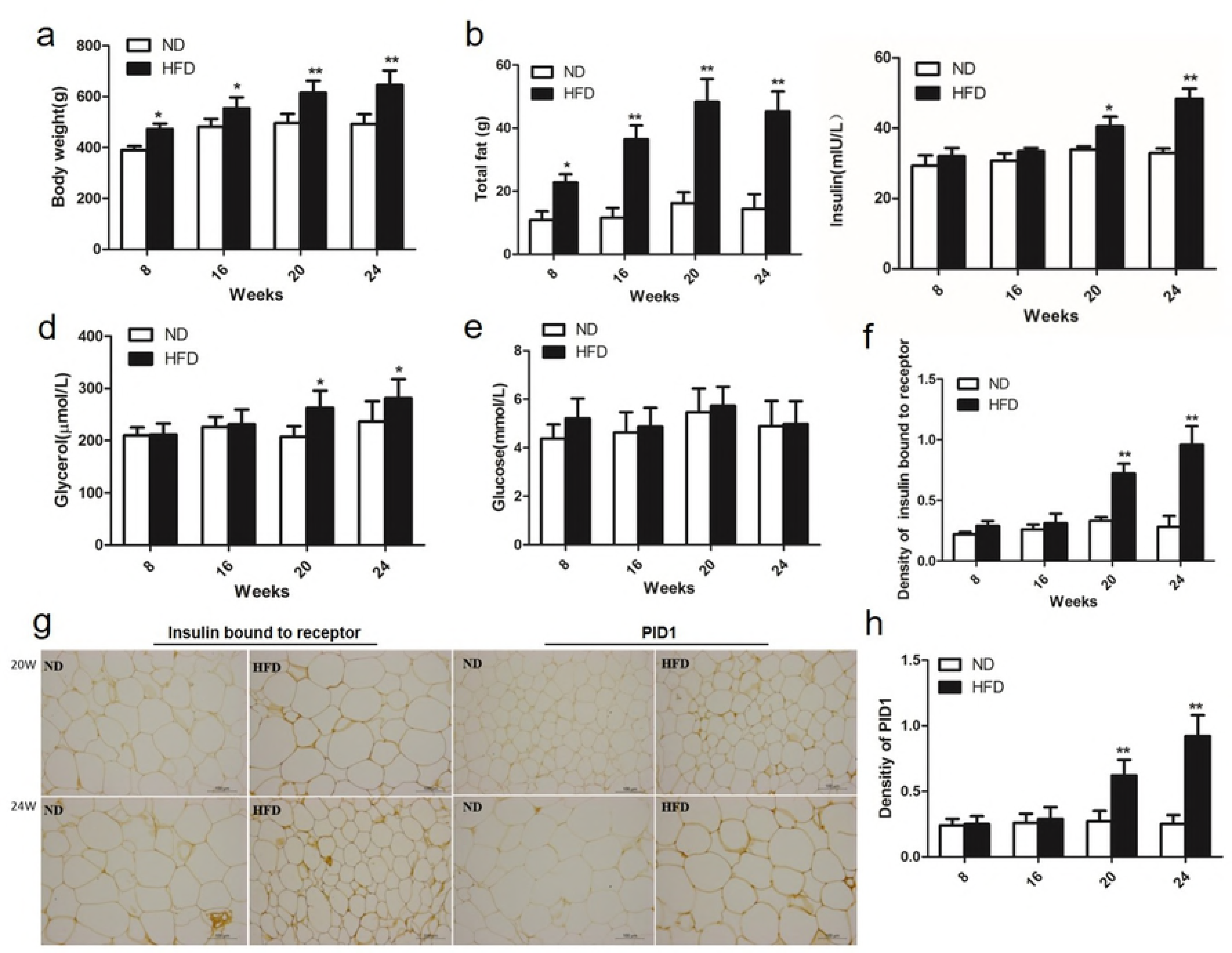
Measures of body weight, total fat, insulin, glycerol and blood glucose of Sprague-Dawley rats fed either a chow or high-fat diet at 8, 16, 20, and 24weeks, and detection the expression of insulin bound to receptor and PID1 in WAT. (a)Body weight. (b)Total fat. (c)Plasma levels of insulin. (d) Plasma levels of glycerol. (e) Plasma levels of glucose. (f) Density of insulin bound to the receptor. (g) Immunostaining of insulin bound to receptor and PID1 in WAT. (h) Density of PID1. Values represent mean ±SE, * P<0.05 compared with ND; ^**^P<0.01 compared with ND.

### Serum glucose, insulin and glycerol levels in HFD and ND groups

After 20 weeks of the diet, the glycerol values for the HFD groups were significantly increased compared with ND groups (Figure 1d). The same differences for insulin levels were also observed after 20 weeks of the diet, with a further increase in high fat diet feeding period (Figure 1c), whereas there were no significant differences in serum glucose leaves throughout the 24-week dietary period (Figure 1e), suggesting a difference in the metabolic response to the diet between the 2groups. Moreover, there was a good correlation in all 2 groups between insulin and glycerol (r=0.57, P<0.05). Since glycerol was an indicator reflecting the lipolysis, we hypothesized that antilipolytic effect of insulin was impaired in HFD rats.

### The levels of Insulin binding to white adipose tissue (WAT)

After 20 weeks of HFD feeding, serum insulin levels of HFD groups was significantly increased compared with ND groups. To assess weather high blood insulin in HFD groupshave an effect on insulinreceptor of adipose tissue. We further measured the levels of insulin bound to receptor of adipose tissuein 2 groups of rats by usingimmunohistologyat8, 16, 20, and 24 weeks. The results showedno differences among 2 groups after 8 and 16 weeksof the diet. However, after20and 24 weeks of HFD feeding, insulin bound to the receptorwasrespectively increased 2.1-fold and 3.8-foldinHFD groupscompared with ND groups (Figure1g-f). The result parallels the increase in blood insulin observed in HFD groups versus the ND groups.

### Expression of PID1 in WAT

We also examined whether the expression of PID1 were altered in WAT of HFD groups compared with ND groups. After 20 weeks of the diet, protein expression of PID1 exhibited a significantly increased in HFD groups compared with ND groups. After 24 weeks of the HFD feeding, PID1expression in HFD groups showed a further increased compared with those on the 20-week diet and was significantly higher than that for ND groups (Figure1g, h), Furthermore, there were positive correlations between PID1 and glycerol levels in2 groups (r=0.42, P<0.05), suggesting that PID1 may play a role in lipolysis.

### Insulin suppresses glycerol release in 3T3-L1via activation of Akt signaling Pathway

To study the effect of insulin on lipid metabolism using 3T3-L1 cell lines. After 3T3-L1cells were fully differentiated, we started off examining dose-dependent effects of insulin in adipocyte lipolysis. Inaccordance with previous studies [11], we found that insulin produced a concentration dependent decrease in glycerol release with significant elevation detectable at 100 nM insulin (Figure 2a, d), and the intracellular lipids increased as insulin dose increased.

**Figure 2.**
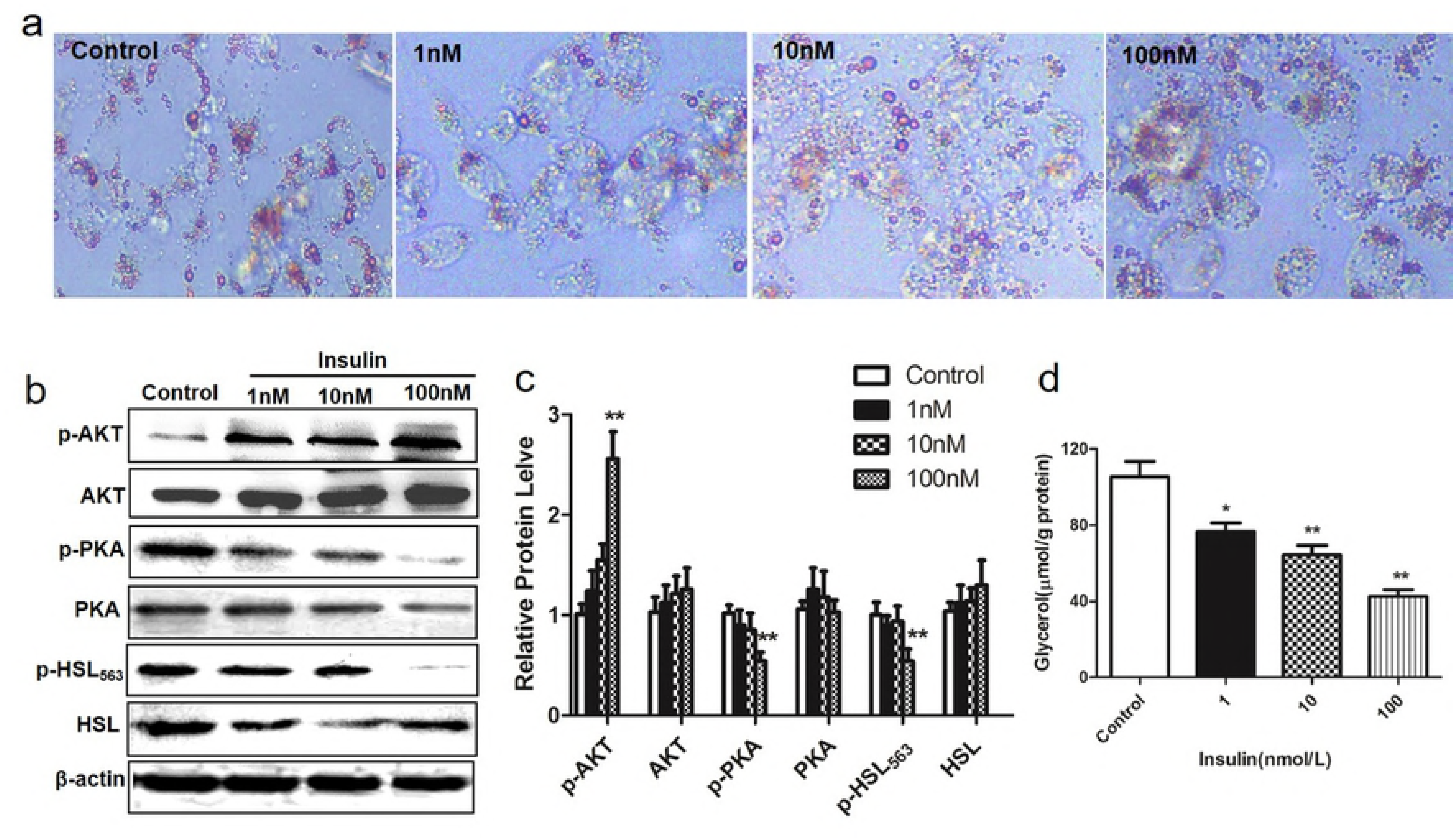
Insulin inhibits lipolysis via AKT/PKA/HSL pathway in 3T3-L1 adipocytes. Differentiated 3T3-L1 adipocytes were treated with insulin at different doses (1-100nM). (a) Cellular triglycerides were measured by oil red O staining.(b-c) Protein expression of AKT, phosphorylated AKT (P-AKT), PKA, phosphorylated PKA (P-PKA), HSL and phosphorylated HSL (P-HSL) were determined by western blot. (d)Glycerol released into medium. ^*^P<0.05 vs. untreated cells; ^**^P<0.01 vs. untreated cells.

We also detected whether Akt were required for insulin’s suppression of lipolysis. when adipocytes were fully differentiated, cells were treat with 1, 10 and 100nM insulin for 24 h. As expected, insulin dose-dependently induced phosphorylation of AKT in 3T3-L1 cell lines. As insulin significantly decreased glycerol release and activated AKT expression at the concentration of 100nM (Figure 2a,b), we used this concentration in the following experiments. Differentiated adipocytes were transfected with AKT siRNA, incubated in the presence of 100nMinsulin. AKT siRNA transfection led to>80% knockdown of the target genes and increased glycerol release in 3T3-L1 (Figure 3b,c). These results suggest that antilipolytic effect of insulin is mediated by AKT in 3T3-L1 adipocytes.

### Effect of activation on the insulin induced decrease in PKA and HSL

Because the current view holds that insulin signaling inhibits lipolysis by reducing PKA and HSL activity [18]. Firstly, we assessed weather insulin inhibits lipolysis via affected the phosphorylation of PKA and known PKA substrates. Secondly, we assessed how siRNA knockdown of AKT or treatment with PKA inhibitors affected the phosphorylation of PKA and HSL. After the addition of different doses of insulin, we analyzed the phosphorylation of HSL at its major PKA site and observed the phosphorylation of PKA. We observed that PKA and HSL phosphorylation levels were significantly reduced on insulin exposure (Figure 2b, c). On the other hand, the AKT siRNA partially reversed the inhibition of PKA and HSL phosphorylation by insulin treatment (Figure 3a,b). Western blot analysis showed that PKA inhibitors pretreatment also dose-dependently reversed the inhibitory effect of insulin on HSL and glycerol release in 3T3-L1(Figure 3d, e). These data confirm that AKT activated by insulin can results in phosphorylation of PKA and HSLand suppresses glycerol release in 3T3-L1.

**Figure 3.**
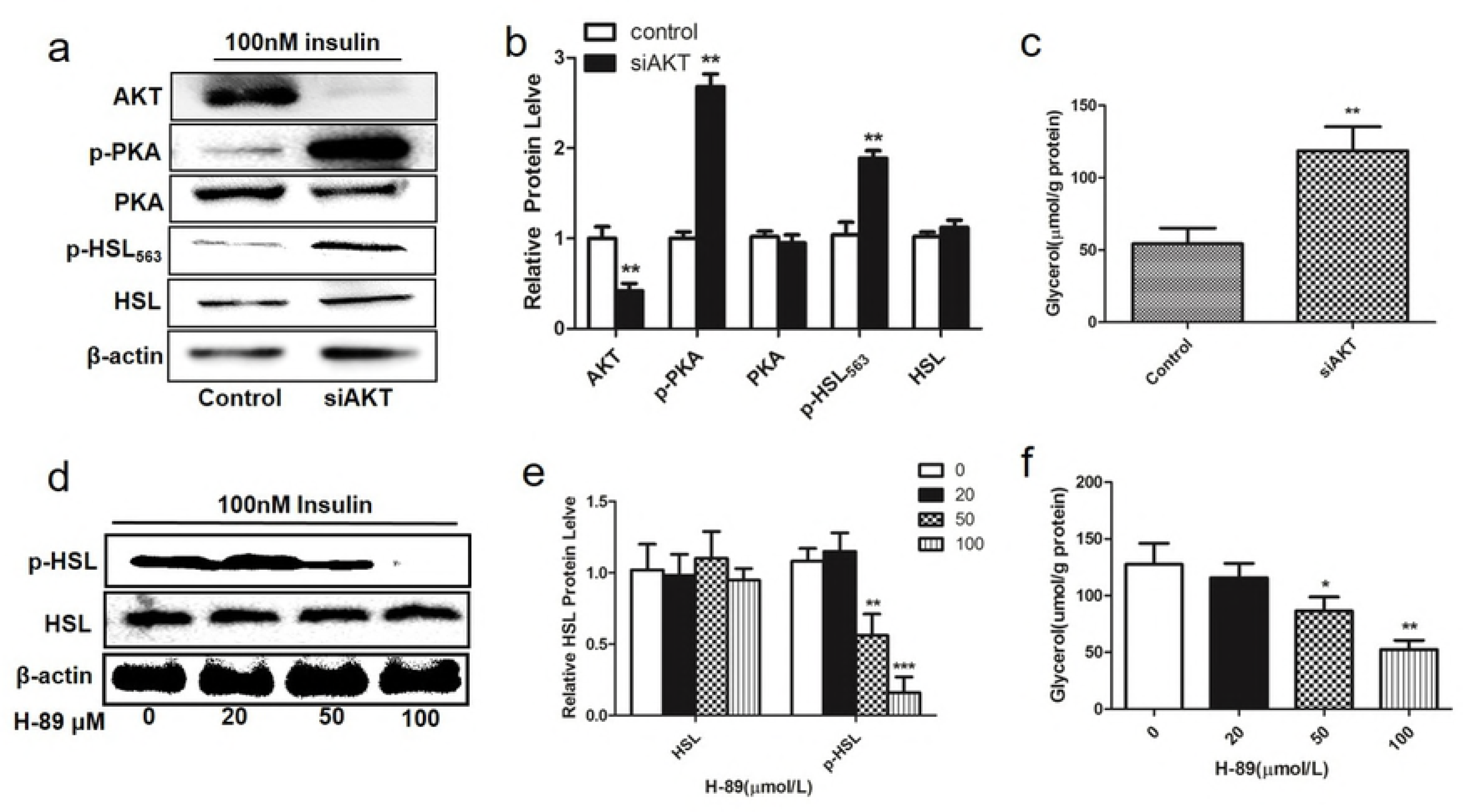
The effects of depletion of AKT or PKA on insulin regulates lipolysis. Differentiated adipocytes were transfected with AKT siRNAortreated with PKA inhibitor, incubated in the presence of 100nMinsulin. (a) Effects of AKT siRNA on insulin induced PKA and HSL protein activation. (b) Relative protein expression of AKT, PKA, phosphorylated PKA (P-PKA), HSL and phosphorylated HSL transfected with AKT siRNA in 3T3-L1 adipocytes. (c) Glycerol released into medium after transfecting with siRNA targeting AKT. (d) Effects of PKA inhibitors on insulin induced HSL protein reduction. (e) Relative protein expression of HSL and phosphorylated HSL treated with PKA inhibitor in 3T3-L1 adipocytes. (f) Glycerol released into medium after treating with PKA inhibitor. ^*^P<0.05; ^**^P<0.01.

### Effects of PID1 over-expression on lipolysis and phosphorylation of AKT/PKA/HSL signaling molecules

To understand the underlying mechanisms by which PID1 affected lipid metabolism, we investigated the effect of PID1 on lipolysis and the proteins involved in insulin signaling inhibits lipolysis. Differentiated adipocytes were transfected with PID1 plasmids, incubated in the presence of 100nM insulin. And then we examined the effects of PID1 over-expression on thelipolysisin response to insulin. Glycerol release was approximately 40% higher in PID1-overexpressing cells than in control cells(Figure 4e). Additionally, PID1 over-expression resulted in noticeable inhibition of insulin-induced Akt serine phosphorylation(Figure 4 c-d). We also evaluated thephosphorylation of the PKA and HSL, which are downstream signaling molecules of the AKT in insulin antilipolytic signaling pathway. We found that PKA and HSL phosphorylation levels were significantly increased in PID1 over-expression cells (Figure 4c-d). This indicates that PID1 inhibits insulin antlipolytic signaling pathway, which involves the inhibition of AKT phosphorylation.

**Figure 4.**
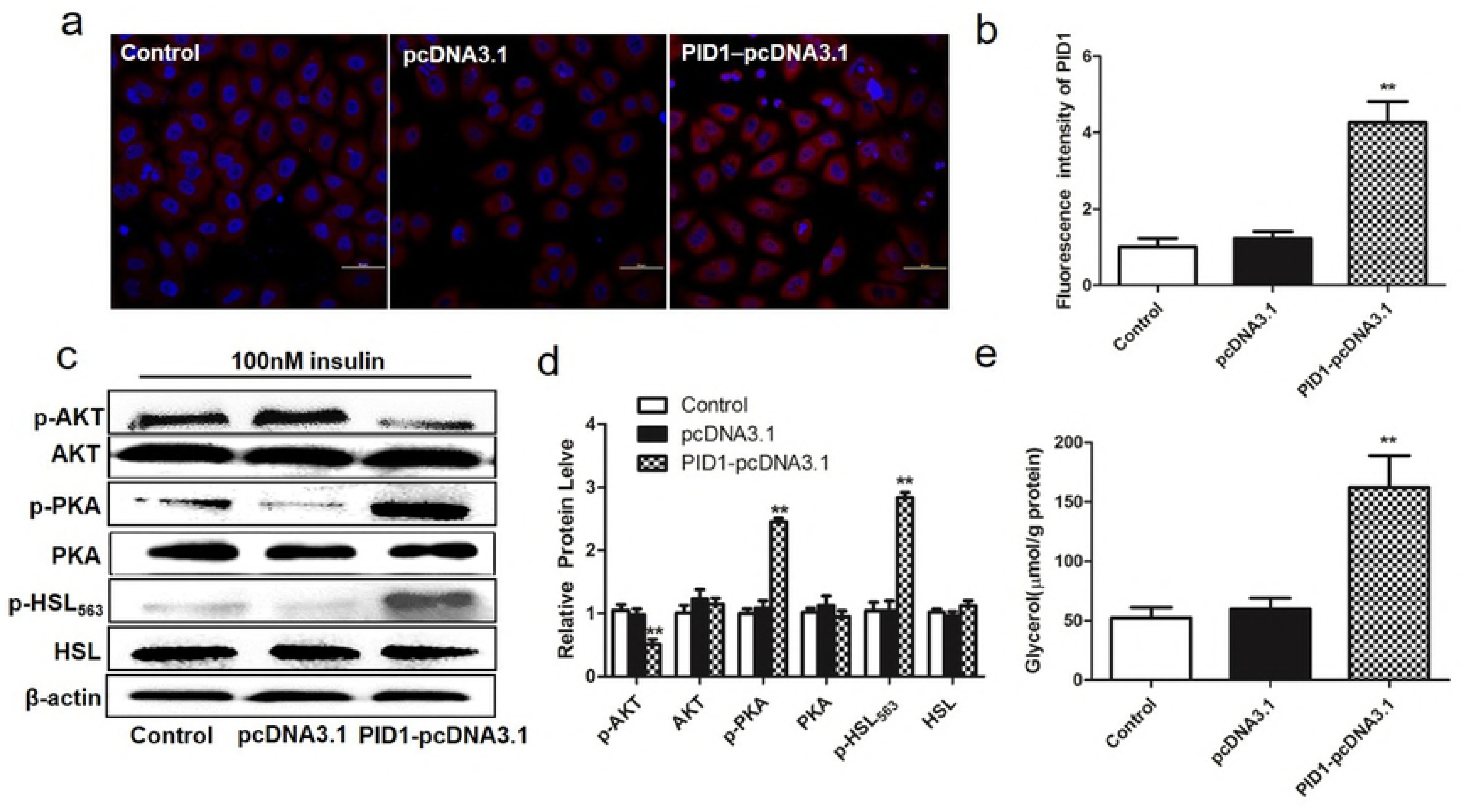
Effects of PID1 over-expression on lipolysis and phosphorylation of AKT/PKA/HSL signaling molecules. Differentiated adipocytes were transfected with PID1 plasmids, incubated in the presence of 100nM insulin.(a-b)Immunofluorescence analysis was performed to confirm the expression of PID1 gene in empty vectors cells, PID1-overexpressing cells, and control cells. (c-d) Effects of PID1overexpression on AKT, phosphorylated AKT (P-AKT), phosphorylated PKA (P-PKA), HSL and phosphorylated HSL protein expression. (e) Effects of PID1 over expression on lipolysis. ^*^P<0.05; ^**^P<0.01.

## Discussion

Our animal experiments show that PID1 expression is increased in adipose tissue from HFD rat and positive correlation with insulin levels and lipolysis. Mechanistically, in 3T3-L1 adipocytes, we found that antilipolytic effect of insulin is mediated by AKT and AKT activated by insulin can results in phosphorylation of PKA and HSL and suppresses glycerol release. However, over-expression of PID1, via inhibiting of insulin-induced activation of AKT, leads to activating phosphorylation of PKA/HSL cascade and promotes lipolysis.

The levels of circulating FFA depend primarily on the rates of lipolysis in the adipose tissue. One of the key physiological functions of insulin as the major anabolic hormone in the body is to restrain lipolysis and to promote fat storage in adipose tissue in the postprandial state[12, 13]. In vitro experiments, we found that insulin activated AKT expression levels in dose-dependent manner, in parallel with decreased glycerol release and increased lipid droplet size. PKA/HSL cascade phosphorylates a wealth of proteins to exert variousbiological functions in different cell types. In adipocytes, this pathway mediates lipolytic effects of several hormones[14, 15]. We showed that AKT depletion activated PKA and HSL phosphorylation and ameliorated the inhibitory action of insulin on lipolysis. PKA inhibitors pretreatment also dose-dependently reversed the inhibitory effect of insulin on HSL and glycerol release. Our data with regard to the mechanism by which insulin inhibits lipolysis are consistent with previous studies, which showed that the inhibitory effect of insulin on lipolysis is attributed primarily to the inhibition of cAMP-mediated signaling to HSL via Akt-dependent [16, 17]. There are considerable data implicating a defect in antilipolysis as a critical etiological abnormality initiating the positive amplifying circuit that characterizes insulin resistance [18, 19]. However, the molecular mechanism by which impaired control of lipolysisin obesity is still unknown at this time. PID1 which contains a phosphotyrosine binding (PTB) domain were subtracted from normal-weight subjects using suppression subtractive hybridization (SSH)[20–22]. The PTB domain usually binds to phosphorylated tyrosine residues and functions in signal transductionby growth factor receptors [23]. In adipocytes and musclecells, PID1 also inhibits insulin-mediated phosphorylationof IRS-1, and insulin-mediated translocation of the GLUT-4 glucose transporter to the membrane, resulting indecreased glucose uptake [24, 25]. Therefore, we hypothesize that the PTB domain of PID1 might impair tyrosine phosphorylation of insulin signaling molecules that inhibit lipolysis. In vivo experiments, PID1 expression is increased in adipose tissue from HFD rat and positive correlation with insulin levels and lipolysis. Consistently, other studies also showed that adipose PID1 expression increased in obesity [26]. Meanwhile, we noticed that over-expression of PID1 significantly increased lipolysis in 3T3-L1 cells. To further investigate the molecular mechanism by which PID1 increases lipolysis, we examined the levels and phosphorylation of proteins involved in insulin signaling for lipid metabolism. The results showed that PID1 decreased the insulin-stimulated serine phosphorylation of Akt, and PKA and HSL phosphorylation levels were significantly increased in PID1 over-expression cells. Based on the seresults, we concluded that over-expression of PID1 promotes lipolysis in 3T3-L1 adipocytes mainly through blocking the AKT/PKA /HSL insulin pathway.

## Conclusion

In conclusion, our results demonstrate that PID1 over-expression promotes lipolysis invitro by attenuating the AKT/PKA /HSL insulin pathway. From in vivo observation to molecular mechanism studies, our results show that activation of PID1 in adipose tissue increases lipolysis by altering the antilipolytic action of insulin. It is believed that impaired control of lipolysis in obesity, which increases circulating FFAs, leads to systemic insulin resistance. Therefore, elucidating the mechanism of PID1-induced lipolysis may constitute a new strategy to ameliorate adipocyte lipolysis and hence to improve insulin sensitivity.

## Funding statement

This work was supported by the Nature Science Foundation of China No. 81172689.

## Conflict of interest

No potential conflicts of interest relevant to this article were reported.

